# Independent control over cell patterning and adhesion on hydrogel substrates for tissue interface mechanobiology

**DOI:** 10.1101/2022.11.16.516785

**Authors:** Louis S. Prahl, Catherine M. Porter, Jiageng Liu, John M. Viola, Alex J. Hughes

## Abstract

Replicating organizational principles that establish fine-scale tissue structure is critical to our capacity for building functional replacement tissues. Tissue boundaries such as epithelial-mesenchymal interfaces are engines for morphogenesis *in vivo*. However, despite a wealth of micropatterning approaches available to control tissue size, shape, and mechanical environment *in vitro*, fine-scale spatial control of cell composition within tissue constructs remains an engineering challenge. To address this, we augment DNA “velcro” technology for selective patterning of ssDNA-labeled cells with long-term culture on mechanically defined polyacrylamide hydrogels. We co-functionalize photoactive benzophenone-containing polyacrylamide gels (BP-PA gels) with spatially precise ssDNA features that confer temporary cell adhesion and with extracellular matrix (ECM) proteins that confer long-term adhesion. We find that co-functionalization does not compromise ssDNA patterning fidelity or cell capture, nor hydrogel mechanical properties or mechanosensitive fibroblast spreading, enabling mechanobiology studies of precise cell interfaces. We then co-pattern colonies of fibroblasts and epithelial cells to study interface formation and extracellular signal-related kinase (ERK) activity at cellular contacts. Combining DNA velcro and ECM functionalization approaches provides independent control of initial cell placement, adhesion, and mechanical environment, constituting a new tool for studying biological interfaces and for programming multicellular interactions in engineered tissues.

## Introduction

Tissues often consist of multiple cell populations with fine-scale spatial organization that is tightly linked to their biological function. These vary from specialized cellular niches that persist throughout adult life such as intestinal villi (Noah et al., 2011) and hair follicles (Millar, 2002) to transitional patterns within a single cell population, such as the differentiation of germ layers in the early embryo (Solnica-Krezel and Sepich, 2012). Cell sorting and boundary formation mechanisms establish and enforce this hierarchical organization. Epithelial-mesenchymal interfaces are a form of heterotypic tissue boundary driving vertebrate embryonic development, for example in tooth cusp formation (Dassule and McMahon, 1998; Mammoto et al., 2011) pharyngeal cartilage formation (Talbot et al., 2016), and branching morphogenesis in the lung, salivary gland, and kidney (Lang et al., 2021; Varner and Nelson, 2014). Disrupted epithelial-mesenchymal interactions can cause structural defects that ultimately impair adult organ function (Müller et al., 1997). Barrier breakdown also presages dissemination of epithelial-derived tumor cells into healthy tissue (Provenzano et al., 2006). The spatial complexity of such interfaces *in vivo* demands that advanced co-culture systems created to reconstitute them *in vitro* have controlled cellular composition and geometry. Such systems would be particularly useful in organizing tissues for regenerative medicine or as models of cellular interactions in development and disease.

Recapitulating fine-scale tissue patterning at relevant length scales *in vitro* is a current challenge in tissue engineering. Micro-stencils and ‘wound healing’ tissue culture inserts loaded with multiple cell populations produce simple and reproducible tissue interfaces (Brayford et al., 2019; Heinrich et al., 2022), but these are low throughput and take hours for cells to close the intervening gap between cell populations. Other microstencil-based hierarchical patterning approaches can provide finer-scale hierarchical tissue patterning (∼100 μm) but involve stepwise surface passivation and de-blocking protocols between patterning different cell types (Javaherian et al., 2014, 2015) or are limited to a single patterned cell type and a second introduced as backfill cells (Bhatia et al., 1998; Shen et al., 2014). Microcontact printing or UV photopatterning approaches reliably deposit extracellular matrix (ECM) protein features that enforce geometric constraints upon tissues down to the single cell scale (∼10 μm) (Jimenez et al., 2021). Such approaches have been successfully implemented on glass, hydrogel, and elastomeric substrates (Liu et al., 2022; Smith et al., 2018; Tang et al., 2012). However, since the initial cell patterning and eventual adhesion footprints are one and the same, micro-printed ECM substrates are limited in their utility to assays in which tissues are restricted in their migration after patterning. Other photolithographic methods such as digital micromirror device projection or direct laser writing afford high spatial resolution and dynamic control over protein patterning (Missirlis et al., 2022; Strale et al., 2016). These techniques permit hierarchical cell co-patterning by dynamically introducing new ECM ligands through biotin-avidin interactions (Strale et al., 2016), yet remain limited by a lack of tools to engineer patterned cell deposition independently of ECM adhesive properties.

DNA-programmed assembly of cells (DPAC) overcomes some of these limitations by patterning cells using single-stranded DNA (ssDNA) deposits as temporary adhesion ligands. Target cells are labeled with lipid-modified complementary ssDNAs and captured at the surface by Watson-Crick-Franklin base pairing (Scheideler et al., 2020; Todhunter et al., 2015). ssDNA deposits can be reliably printed at tissue-relevant spatial scales (∼10-100 μm) and cell patterning can be multiplexed, using a unique ssDNA sequence to place each cell type. DPAC approaches have previously relied on aldehyde-coated glass slides to support covalent linkage of ssDNA to the substrate. While this allows for subsequent cell adhesion to the glass or release and transfer into an encapsulating hydrogel overlay, reliance on a glass substrate limits other opportunities facilitated by compliant hydrogels, such as measurement of cell mechanical forces (Pelham and Wang, 1997) or influencing differentiation through substrate stiffness (Engler et al., 2006). We recently published a DPAC approach involving rapid photolithographic patterning of ssDNAs onto polyacrylamide hydrogels containing a benzophenone-methacrylate (BPMAC) co-monomer (Viola et al., 2020). BPMAC is photoactive, creating covalent bonds with polypeptides and ssDNA oligomers upon irradiation with 250-365 nm light (Dormán and Prestwich, 1994; Hughes and Herr, 2012; Viola et al., 2020). However, these benzophenone-polyacrylamide (BP-PA) hydrogels are not inherently cell-adhesive, presenting an opportunity to control cell adhesion by orthogonal means.

Here, we present a multi-step procedure for fabricating BP-PA hydrogel substrates, photopatterning ssDNA, and ECM functionalization. These substrates enable studies of non-autonomous cell interactions across heterotypic cell interfaces in a convenient 2D format. We find that ssDNA photolithography is orthogonal to established polyacrylamide surface chemistry approaches that enable cell adhesion through surface-bound ECM proteins, meaning that cell patterning and cell adhesion chemistry can be independently controlled. We demonstrate that ssDNA photocapture and UV exposure do not disrupt BP-PA hydrogel mechanics or fibroblast mechanosensing, facilitating mechanobiology studies. Photopatterning multiple ssDNA sequences allows us to rapidly produce colonies of multiple cell types with defined initial size, shape, position, and boundary geometries, allowing cells to dynamically interact in space and time. Finally, we demonstrate that the boundary geometry alone influences cytoskeletal arrangements and extracellular signal-related kinase (ERK) activity at interfaces within micropatterned composite epithelial-mesenchymal tissues. This multiplexed patterning approach enables new possibilities for spatially resolved studies of the mechanobiology of cell and tissue interactions. This has potential impacts in broad ranging fields from regenerative medicine and synthetic biology to disease modeling.

## Results

### BP-PA hydrogels support a combined cell patterning and long-term 2D tissue culture strategy

Augmenting the already versatile polyacrylamide hydrogel system with DNA-patterned cell adhesion would realize a significant performance benefit to the production and study of cell behavior at tissue interfaces. We therefore established a multi-step process for photopatterning ssDNA, surface functionalization with ECM, and cell patterning onto BP-PA hydrogels. Quartz-chrome photomasks enable simultaneous deposition of up to 10^7^ ssDNA features (Viola et al., 2020) (∼10-100 μm feature size) and/or polypeptides (**figure S1**) through a UV photochemical reaction with BPMAC co-monomers in the hydrogel (**figure 1a**). Photopatterned ssDNA features then define capture sites for lipid-ssDNA labeled cells, through base pairing. Washing the hydrogel removes excess cells, leaving cell patterns anchored to ssDNA features (**figure 1b**). In the present study, we used an “open face” format using a commercially available 8-well chambered slide that facilitates subsequent long-term culture in a convenient format for imaging (see: **materials & methods**).

**Figure 1.**
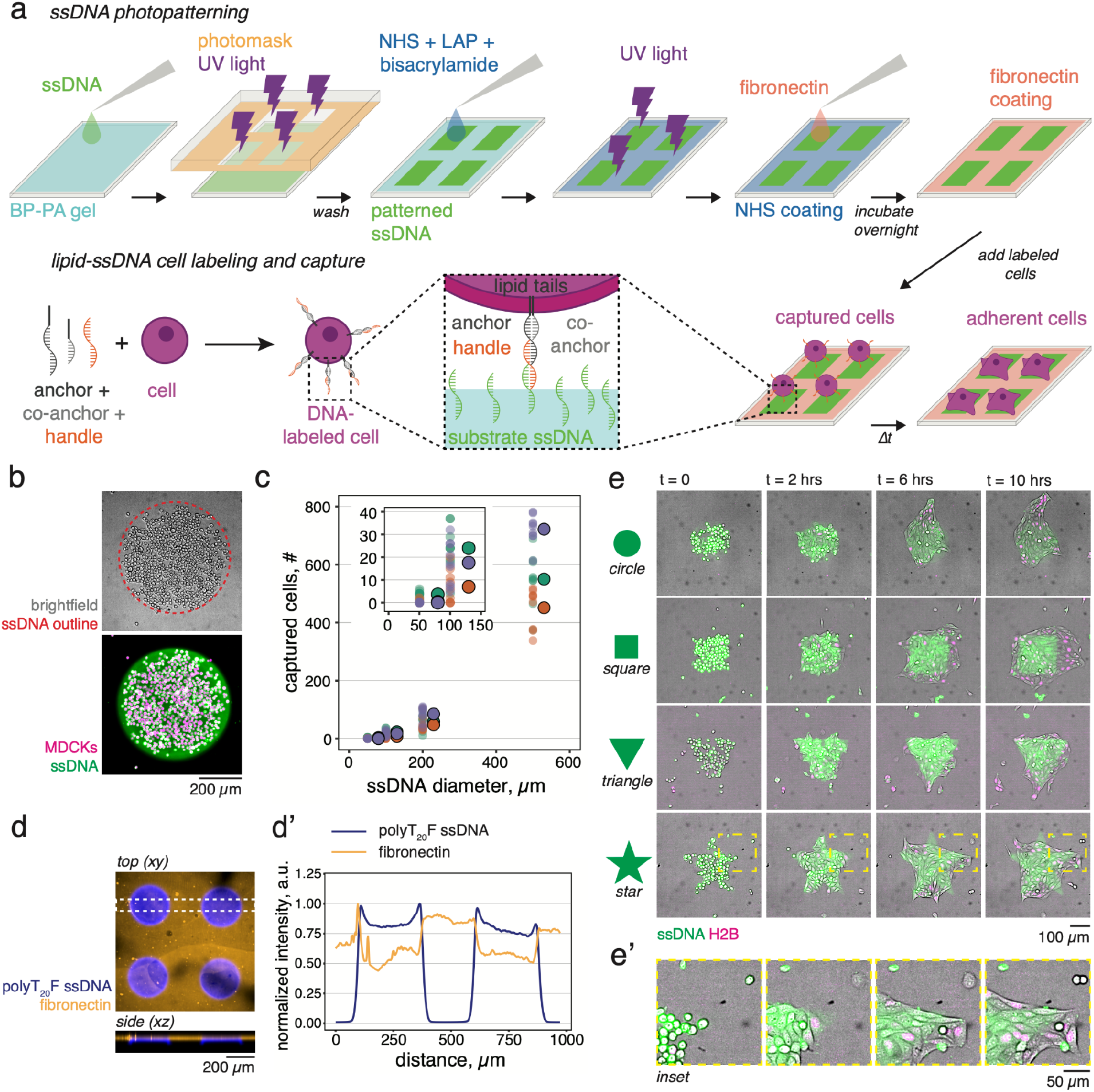
Cell capture and long-term adhesion on polyacrylamide substrates through orthogonal ssDNA and ECM patterning. **a**. Schematic of the *top*, ssDNA photolithography and BP-PA gel surface functionalization with fibronectin, and *bottom*, capture of lipid-ssDNA labeled cells. Cell labeling with lipid-ssDNA and sequence-matched “handle” ssDNA enables specific cell capture to hydrogel-bound ssDNA patterns. Captured cells subsequently adhere to the substrate using fibronectin ligands. **b**. Brightfield (*top*) and fluorescence (*bottom*) images of MDCK cells captured on a 500 μm diameter circular ssDNA feature. **c**. Quantification of MDCK cell capture (#) on 50, 100, 200, and 500 μm diameter ssDNA circles. Data depict *n* = 9 features per ssDNA diameter per three independent replicate experiments (*n* = 27 features total per ssDNA diameter) and means of each experiment (black borders). Experiment means are shifted +30 μm along the +x axis for clarity. **d**. *Top*, maximum intensity projection (*xy*) of photopatterned ssDNA (polyT_20_F) and fibronectin on a BP-PA hydrogel from a 10x z-stack (2.5 μm per frame, 147.5 μm total height, 59 total frames). *Bottom*, maximum *xz* intensity across the highlighted 25 pixel region. **d’**. Normalized mean fluorescence intensity sampled across the outlined portion of the *xy* image in panel d. **e**. Time lapse image sequences (10 hrs total) of cells adhering to circular, triangular, square, and star-shaped ssDNA features with fixed areas equivalent to a 200 μm diameter circle (A = 3.14×10^4^ μm^2^). **e’** *Inset*, detail of the star-shaped pattern. All experiments were performed on 7.5%/0.25% (Am/Bis) BP-PA hydrogels photopatterned with 200 μM polyT_20_G ssDNA for 90 seconds and functionalized with 20 μg ml^-1^ fibronectin. Cells were labeled with lipid anchors and “G’ handle” ssDNA. Cells in panel **b** are labeled using CellTracker Deep Red and ssDNA in **b** and **e** is visualized using 2x SYBR Gold nucleic acid stain. Photopatterned polyT_20_F ssDNA in panels **d-d’** is visualized with FAM_F’ ssDNA probe and fibronectin is visualized with a rabbit anti-fibronectin primary antibody and Alexa 647 secondary antibodies. See also: **figure S1-S2** and **movie S1**.

To achieve cell capture, we used a previously described lipid-ssDNA universal anchor/co-anchor pair (Viola et al., 2020; Weber et al., 2014) that is compatible with two previously described sense/antisense ssDNA pairs - denoted here as F/F’ and G/G’ (**table s1**). The “F” and “G” sequences contain a string of 20 thymine bases followed by a 20 base pair adhesion sequence (e.g., polyT_20_X_20_). “F’ handle” or “G’ handle” sequences are introduced to cells following tagging with lipid-ssDNA anchor and co-anchor (**figure 1a**). We previously determined that UV exposure times of 60-120 seconds (λ = 254 nm, I = ∼7 mW cm^-2^) in the presence of 200 μM polyT_20_X_20_ ssDNA were sufficient to support cell capture on BP-PA hydrogels (Viola et al., 2020) so we continued to use exposure times within this range. Next, we benchmarked capture efficiency on features of varying size. MDCK cells were efficiently captured on 50-500 μm diameter circular features using the G/G’ ssDNA pair, with larger features predictably capturing more cells (**figure 1d**). However, we note that patterning on smaller features reduces the yield in our “open face” format (only 11/27 or 41% of the 50 μm diameter ssDNA features we measured contained any captured cells, **figure 1c**). Capture efficiency can be improved to single cell resolution using a microfluidic flow cell to introduce and wash cells (Viola et al., 2020). These data show that our open-face assay design achieves similar cell capture properties to previous pDPAC approaches for pattern sizes relevant to the study of tissue interfaces, but in a radically simpler format.

Turning to the goal of combining long-term cell adhesion with patterned cell capture, we established an orthogonal method for functionalizing the BP-PA hydrogel surface with ECM proteins. Providing ECM ligands allows captured cells to adhere to the hydrogel substrate through integrins and other cell adhesion receptors. To functionalize hydrogels with ECM, we modified a previously described photochemical reaction to derivatize acrylamide chains with N-hydroxysuccinimide (NHS) ester groups that couple proteins to the hydrogel surface through primary amines (Lakins et al., 2012). We applied a mixture of a UV photoinitiator compound lithium phenyl-2,4,6-trimethylbenzoylphosphinate (LAP), hydrogel cross linker N,N-methylenebisacrylamide (Bis), and an NHS ester-containing compound (acrylic acid-N-hydroxysuccinimide) and UV exposed the entire hydrogel for 10 minutes (λ = 365 nm, I = 15 mW cm^-2^). After washing, we coupled ECM proteins (20 μg ml^-1^ fibronectin) to the hydrogel surface through overnight incubation (**figure 1d)**. To demonstrate the uniformity of ECM functionalization and ssDNA photopatterning, we probed the fibronectin distribution across ssDNA-patterned and unexposed hydrogel regions by immunofluorescence and confocal imaging. Fibronectin was distributed uniformly across non-exposed hydrogel regions, decreasing marginally in intensity (∼25%) across areas bearing photopatterned ssDNA (**figure 1d’**). This NHS derivatization method gave the best cell adhesion results relative to other commonly used ECM functionalization techniques (e.g., Sulfo-SANPAH), which exhibited dewetting of epithelial layers from previously UV-exposed regions (**Supplemental Note 1, figure S2**).

To test whether ECM functionalization would support long-term cell adhesion, we photopatterned ssDNA features of fixed area (A = 3.14×10^4^ μm^2^, equivalent to a circle with 100 μm radius) but varying shape and introduced lipid-ssDNA-labeled Madin-Darby canine kidney (MDCK) epithelial cells (**figure 1e**). Captured MDCK cells subsequently adhered to the ECM, formed cell-cell junctions, and began to collectively spread as epithelial colonies. Importantly, the intervening ECM functionalization step did not prohibit the capture of lipid-ssDNA labeled cells, nor did ssDNA photopatterning prevent long-term cell adhesion on ssDNA features. We subsequently tested the capacity of our functionalized BP-PA gels to support the adhesion of various cell types following capture on arrays of 250 μm circular ssDNA features, including epithelial cells (MDCK, LLC-PK1), fibroblasts (3T3), and embryonic tissue-derived cells (HEK 293T) (**figure S2c**). After 24 hours, MDCK and LLC-PK1 cells maintained cell-cell contacts and spread as distinct colonies, while 3T3s spread and covered much of the hydrogel surface. HEK 293T cells are a weakly adherent cell line and spread poorly on fibronectin (**figure S2c**), suggesting that some cell types may require optimization of the ECM ligand or concentration to promote long-term attachment. These data together show that augmenting DPAC with ECM functionalization facilitates long-term retention of patterned tissues and time-resolved studies of cell behavior.

### BP-PA hydrogel photopatterning retains gel properties suited to mechanobiology studies

Polyacrylamide gels are desirable for mechanobiology studies because their elastic modulus (*E*) can be tuned by changing the relative amounts of acrylamide (Am) and Bis crosslinker in the pre-polymer solution (Denisin and Pruitt, 2016; Pelham and Wang, 1997). Changes in elastic material properties influence mechanosensitive cell migration, proliferation, differentiation, and intracellular signaling (Bangasser et al., 2017; Engler et al., 2006; Farahani et al., 2021; Solon et al., 2007; Ulrich et al., 2009). To validate that our co-functionalization approach is compatible with these favorable tuning properties, we cast BP-PA hydrogels with varying Am/Bis composition (3-7.5% Am, 0.035-0.25% Bis, **table S2**) and indented them with a 255 μm blunt-ended cylindrical indenter attached to a force sensor (**figure 3a**) (Levental et al., 2010).Polyacrylamide hydrogels can be approximated as linear elastic materials (Anseth et al., 1996; Denisin and Pruitt, 2016) so we were able to directly measure *E* from the time-invariant linear part of force versus indentation depth curves (Levental et al., 2010). To test whether UV-induced ssDNA patterning influenced *E*, we exposed half of each hydrogel in the presence of ssDNA through a quartz microscope slide to permit 254 nm light transmission (+UV) and blocked the other half from UV exposure (ctrl). We measured similar values of *E* for ctrl and +UV hydrogel regions across a range of BP-PA gels cast from 3%-7.5% Am and 0.05%-0.25% Bis (**figure 3b**). Hydrogels cast with 3%/0.05% Am/Bis ratio were ∼14% stiffer on the +UV side (ctrl: 2.4 ± 0.6 kPa, +UV: 2.8 ± 0.7; *n* = 4 hydrogels, *p* = 0.016 by paired t-test), while UV exposure had an insignificant effect on *E* in all other cases. These data validate that BP-PA gels retain tunable elastic material properties following ssDNA photopatterning.

**Figure 2.**
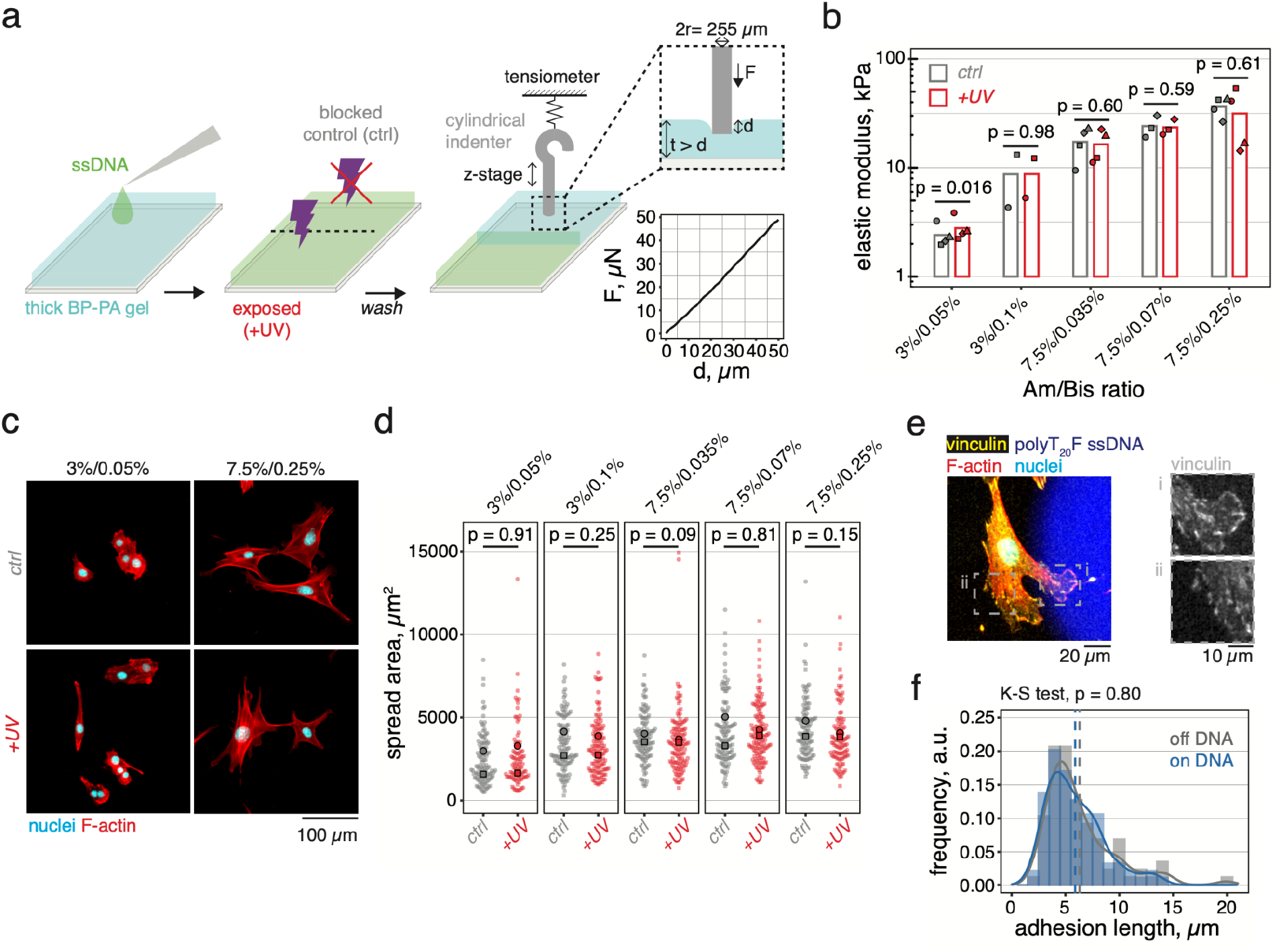
Photopatterned BP-PA hydrogel mechanics, fibroblast spreading, and focal adhesion formation are comparable to control BP-PA gels. **a**. Schematic of hydrogel UV exposure and microindentation with example force vs. indentation depth curve obtained for a 255 μm diameter cylindrical indenter on a 3%/0.05% Am/Bis ratio BP-PA hydrogel. Hydrogels were incubated in ssDNA (200 μM polyT_20_G) and exposed to 254 nm light for 90 seconds through one half of a quartz slide (+UV), with the other side blocked from UV exposure (control). **b**. Quantification of *E* for BP-PA hydrogels cast with 3-7.5% Am, 0.01-0.25% Bis, and 3 mM BPMAC (see **table S2** for mean ± s.d. for *n* = 2-4 hydrogels per Am/Bis composition). Individual data points are identified by shape, bar heights represent overall mean. **c**. 3T3 fibroblasts spreading on +UV and control hydrogel regions functionalized with 20 μg ml^-1^ fibronectin. Cells adhered for 16-24 hours prior to fixation and staining for F-actin and nuclei (DAPI). **d**. Quantification of 3T3 spread area on control and +UV hydrogel regions from BP-PA hydrogels cast with varying Am/Bis compositions; control, *n* = 108, 117, 116, 119, 110 cells and +UV, *n* = 92, 128, 129, 135, 110 cells. Data are pooled from two independent replicate experiments identified by marker shape, individual experiment means are overlaid onto distributions (black borders), *p*-value comparisons between groups are from two-sided Kolmogorov-Smirnov tests. **e**. 3T3 fibroblast adhered to a 7.5%/0.25% (Am/Bis) BP-PA hydrogel containing 250 μm circular ssDNA features (labeled with FAM_F’), nuclei (DAPI), F-actin, and focal adhesions (vinculin). *Insets*, detail on regions marked by dashed boxes. **f**. Quantification of *L*_*adhesion*_ for cells on unexposed (off DNA) and UV-exposed (on DNA) hydrogel regions. Vertical dashed lines represent the mean of *n* = 94 (off DNA) and 87 (on DNA) focal adhesions measured from *n* = 29 cells pooled from two independent experiments (biological replicates). *p*-value was computed using a two-sided Kolmogorov-Smirnov test.

**Figure 3.**
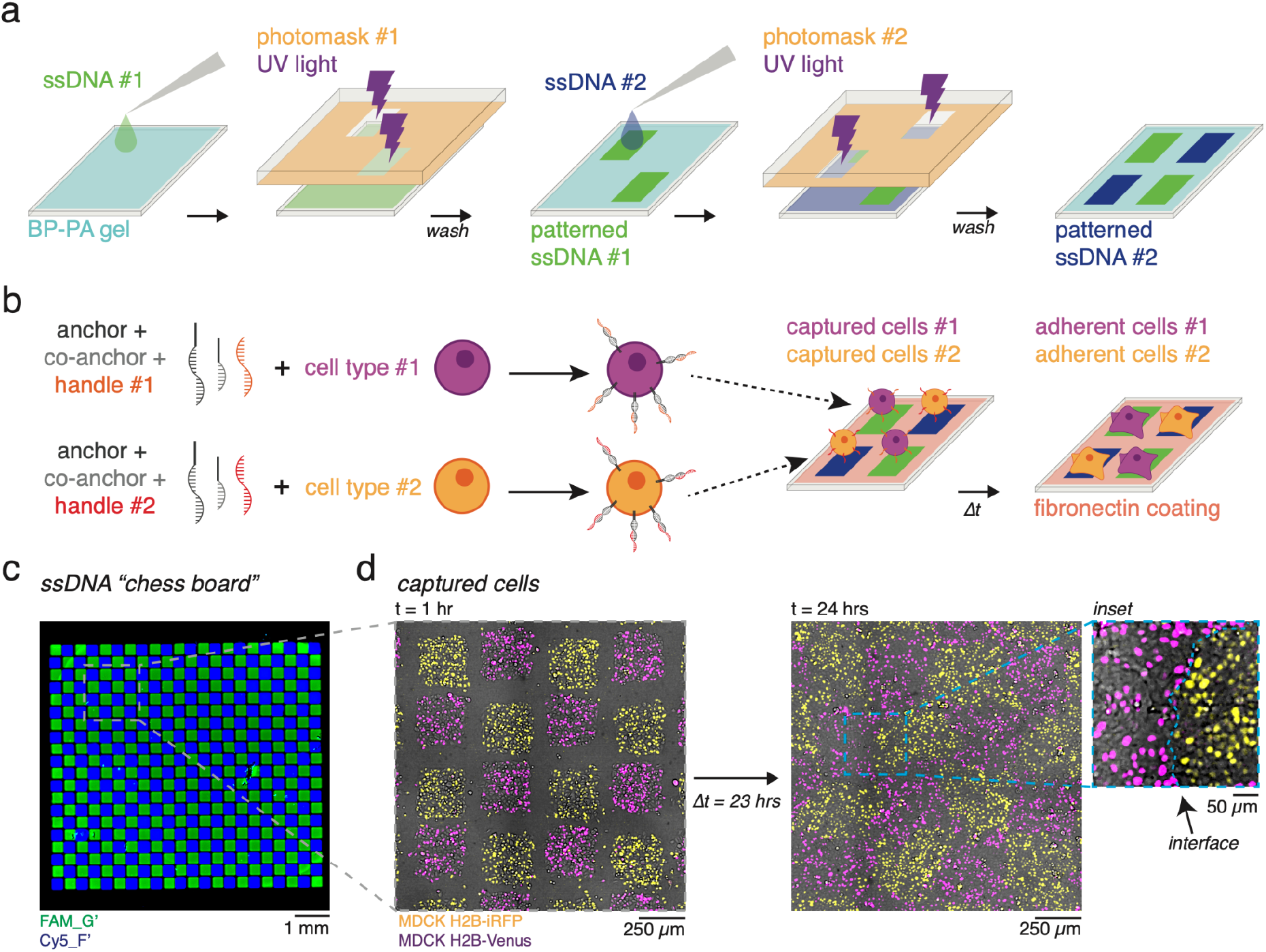
ssDNA multiplexing supports production and long-term adhesion of patterned tissues. **a**. Schematic of iterative ssDNA photopatterning. **b**. Schematic for lipid-ssDNA labeling and patterning of multiple cell populations. Note surface fibronectin functionalization is not shown schematically, but is performed before cell capture. **c**. Confocal micrograph of polyT_20_F and polyT_20_G sequences photopatterned into a chess board pattern of alternating squares (250 μm side length) across an entire culture well. ssDNAs were sequentially patterned onto a 7.5%/0.25% (Am/Bis) BP-PA hydrogel by 254 nm UV, t = 60 s exposure for each ssDNA. Hydrogels were also functionalized with 20 μg ml^-1^ fibronectin. ssDNA patterns are visualized with 1 μM FAM_G’ and 1μM Cy5_F’ fluorescent ssDNA probes. **d**. Two populations of MDCKs expressing fluorescent H2B constructs captured on a similar chess board pattern. Cells were imaged t = 1 hr after capture and at t = 24 hrs. MDCK H2B-Venus cells were patterned using the G/G’ ssDNA pair and MDCK H2B-iRFP cells were patterned using the F/F’ ssDNA pair (**table S1**). *Inset*, detailed view of interface. Data are representative of two independent experiments. See also: **figure S3**.

We next asked whether photopatterning of BP-PA hydrogel surfaces influenced cell-hydrogel interactions. Fibroblast spreading increases with increasing *E* within a physiologically relevant range (Solon et al., 2007). Adherent 3T3 fibroblasts showed similar spread area vs. *E* relationships on the control and +UV regions of BP-PA hydrogels across each Am/Bis composition that we tested (**figure 3c-d, table S2**). We also characterized the formation of focal adhesions, which are protein complexes that enable cell interpretation of substrate mechanical properties and ECM ligand density (Peyton and Putnam, 2005). To test whether focal adhesion formation differed between unexposed and photopatterned hydrogels, we let fibroblasts adhere for 16-24 hrs to BP-PA hydrogels (7.5%/0.25% Am/Bis ratio, **table S2**) functionalized with 20 μg ml^-1^ fibronectin and bearing arrays of 250 μm circular photopatterned ssDNA features.Immunostaining for the focal adhesion protein vinculin allowed us to measure focal adhesion lengths (*L*_*adhesion*_) on unexposed hydrogel surfaces and FAM_F’ probe-labeled ssDNA features (**figure 3e**). We detected no difference in *L*_*adhesion*_ between adhesions on unexposed hydrogel (*L*_*adhesion*_ = 6.3 ± 3.2 μm, mean ± s.d. of *n* = 94 adhesions) and ones on ssDNA features (*L*_*adhesion*_ = 5.9 ± 2.6 μm, mean ± s.d. of *n* = 86 adhesions) (**figure 3f**). ssDNA photopatterning on BP-PA hydrogels therefore adds a precise cell patterning capability without altering subsequent focal adhesion formation and cell spreading induced by ECM co-patterning (see: **supplementary note 1**).

### Combining multiplexed ssDNA photolithography with ECM functionalization enables production and long-term adhesion of patterned tissue interfaces

Precise spatial control over multiple cell populations would benefit studies of cell contact-related behaviors such as juxtacrine signaling or heterotypic cell adhesion. However, existing micropatterning techniques cannot reproduce complex multicellular patterns or interfaces at tissue-relevant length scale. To overcome this limitation, we took advantage of the minimal cross-reactivity between F/F’ and G/G’ ssDNA pairs during cell capture (Scheideler et al., 2020; Todhunter et al., 2015; Viola et al., 2020) and used them to place multiple cell populations on photopatterned hydrogels. To align and register multiple ssDNA sequences on the same substrate, we adapted an iterative photopatterning process (**figure 3a**), where each successive photomask pattern is registered to fiduciary features from previous steps using a fluorescent ssDNA antisense probe (Todhunter et al., 2015; Viola et al., 2020). We first used this process to create an alternating ‘chess board’ ssDNA pattern and captured two distinct populations of MDCK cells expressing different fluorescent histone 2B (H2B) constructs to label cell nuclei (**figure 3b**). We found no overall difference in mean cell capture efficiency between the two populations (**figure S3**). Patterned cells adhered to surface-bound fibronectin ligands within 1 hr of patterning and spread outward to form cell-cell contacts with neighboring patterned populations, ultimately establishing a confluent monolayer after 24 hrs. Within this time period, cell patterns retained aspects of their initial geometric organization, while showing some boundary evolution through movement and mixing, which occurs even in confluent epithelial monolayers (Javaherian et al., 2014). These data show that serial ssDNA photopatterning steps can be performed without compromising cell patterning fidelity or ECM functionalization, enabling the formation of geometrically controlled tissue interfaces.

### Patterned epithelial-mesenchymal interactions support investigation of physical and biochemical interactions between cell populations

Interacting epithelial and mesenchymal cell populations are crucial to the early formation of a number of embryonic organs (Mammoto et al., 2011; Müller et al., 1997; Talbot et al., 2016), motivating development of advanced culture technologies to accelerate understanding and synthetic construction of epithelial-mesenchymal interfaces. Brayford *et al*. reported that mesenchymal contact inhibition drives sorting of epithelial and mesenchymal cells into distinct populations *in vitro*, as well as the formation of stable boundaries where the two populations interact (Brayford et al., 2019). They also found that boundary stability *in vitro* involves signaling through the Eph/ephrin and extracellular signal-related kinase (ERK) signaling pathways. However, as their experiments used wound healing inserts to create a millimeters-long interface, it is unclear how epithelial-mesenchymal interactions would evolve given different patterning geometries and length-scales.

To test the role of boundary geometry on epithelial-mesenchymal interface formation, we designed two-part photomask patterns that split a 500 μm diameter circle into a concentric circle and annulus of equal area (*r*_*inner*_ = 176 μm, *r*_*outer*_ = 250 μm, *A*_*inner*_ = *A*_*outer*_ = 97264 μm^2^). This design creates composite tissues having one cell type enclosed within another (**figure 4a**). After producing BP-PA hydrogels bearing the two photopatterned ssDNAs and functionalizing them with ECM, we patterned lipid-ssDNA labeled MDCK epithelial cells and 3T3 fibroblasts and used live imaging to study their interactions (**figure 4b**). When fibroblasts were enclosed by a ring of MDCK epithelial cells (“interior mesenchyme” tissues) the inner fibroblast population underwent modest expansion in area outward against the epithelial boundary (**figure 4b-c** and **movie S2**). However, in some cases, fibroblasts exploited discontinuities in the epithelial ring to escape from the interior (**figure S4a**). We also observed a progressive decrease in circularity among interior mesenchyme tissues (**figure 4d**), reflecting increasingly irregular shapes of the fibroblast populations caused by instability of tissue interfaces over time. In the reverse configuration (“interior epithelium” tissues) the collective outward expansion of epithelial cells increased significantly (**figure 4b-c**), leading to compression of fibroblast fields at interfaces.

**Figure 4.**
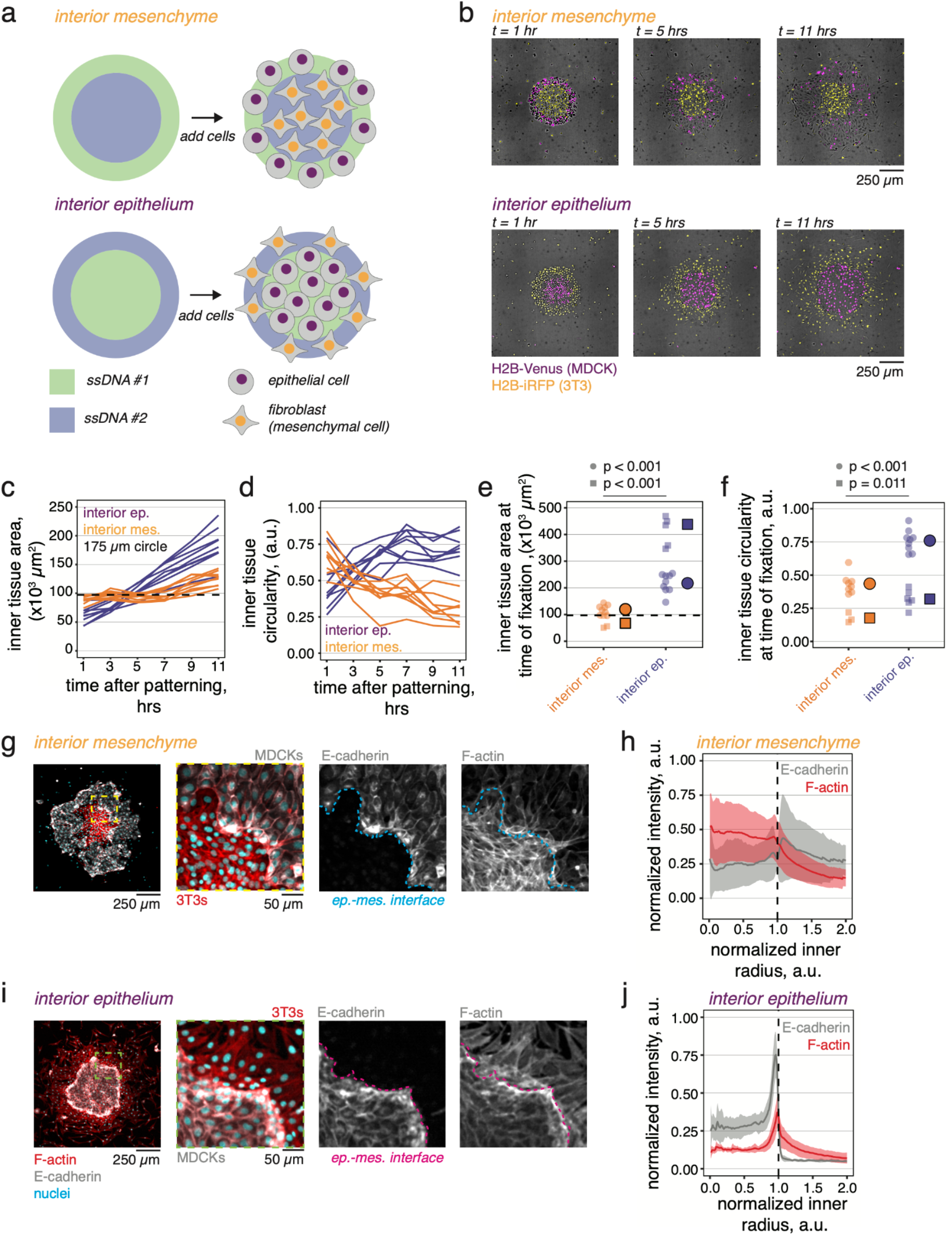
Epithelial-mesenchymal tissue interfaces show emergent organization that depends on initial tissue geometry. **a**. Schematic of two-part ssDNA patterning design consisting of concentric circles. **b**. Example images of co-patterned H2B-iRFP 3T3s and H2B-Venus MDCKs acquired starting 1 hour after patterning and every 2 hours for 10 hours total. **c-d**. Inner tissue area and circularity measured from *n* = 8 interior mesenchyme and *n* = 9 interior epithelium tissues from an example experiment. **e-f**. Summary plots of tissue area and circularity at time of fixation (t = 12-15 hrs after patterning) measured for *n* = 11 interior mesenchyme and *n* = 15 interior epithelium tissues collected from two independent replicates. Individual experiment means (markers with black borders) are offset to the right of each group and data points for each experiment are organized by shape. p-values for each replicate are computed using Welch’s two-sided t-test. Dashed horizontal line in panels **c** and **e** represents the inner tissue patterning radius (r = 175 μm). **g**. Confocal micrograph of an “interior mesenchyme” microtissue stained for F-actin and E-cadherin. *Inset*, composite and single channel images show alignment of epithelial cells and fibroblasts at the interface (dashed lines in each panel). **h**. Radial quantification of normalized E-cadherin and F-actin fluorescence intensity relative to inner tissue radius. **i**. Example “interior epithelium” tissue stained for E-cadherin and F-actin and **j**. radial quantification of normalized fluorescence intensity relative to inner tissue radius. Vertical dashed line marks the tissue interface position (r = 1.0 a.u.), ribbons are mean ± s.d. of tissues analyzed in panel **e**. In all experiments, MDCK cells are patterned using the G/G’ ssDNA pair, while 3T3 cells are patterned using the F/F’ ssDNA pair. Cells were patterned on 7.5%/0.25% (Am/Bis) BP-PA hydrogels sequentially photopatterned with polyT_20_G and polyT_20_F ssDNA (t = 90 s exposure at 254 nm each) and functionalized with 20 μg ml^-1^ fibronectin. See also: **figure S4-S5** and **movies S2-S3**.

Compression increased their local density and alignment at the tissue edge (**movie S3**). As a result, we observed a progressive increase in inner tissue area (**figure 4c**) and circularity (**figure 4d**) across the imaging window, reflecting higher stability of the tissue interface over time. Endpoint measurements of interior tissue area and circularity in culture (after 12-15 hrs) revealed a consistent effect between duplicate experiments (**figure 4e-f**). These data show that tissue compartment area and interface stability depend on patterning boundary conditions, even though the context of cell interaction at the interface is identical at the single-cell scale.

Heterotypic tissue contacts can induce collective alignment of cells and formation of supra-cellular cytoskeletal structures at the interface (Javaherian et al., 2015; Monier et al., 2010). To investigate cytoskeletal organization at the interface, we stained tissues for an epithelial cell marker (E-cadherin) and F-actin. Patterning tissues into a reproducible starting geometry enabled us to quantify the radial intensity of each marker (Dobretsov and Romanovsky, 2006). Fibroblasts located within the center of a composite tissue were densely packed and showed an accumulation of actin stress fibers, leading to an increase in F-actin intensity within the tissue compared to the surrounding epithelium (**figure 4g-h**). By contrast, interior epithelial tissues had a sharp accumulation of actin and E-cadherin at the interface (**figure 4j**). Elongated cell morphologies and alignment perpendicular to the interface (accompanied by accumulation of actin stress fibers within both cell populations, **figure 4i**) suggest elevated tension in the interface cells, which may prohibit cell migration or division across the boundary (Javaherian et al., 2015; Monier et al., 2010). Differences in cell density between colliding tissues may also influence the time evolution of boundary formation, with less dense tissues being displaced by denser ones (Heinrich et al., 2022). Indeed, cell density (using radially-binned average nucleus signal as an approximation) was markedly higher in interior fibroblast tissues compared to their density in the opposite patterning scheme when they bordered epithelial cells (**figure S5**).

These data indicate the importance of boundary geometry in determining equilibrium cell densities and cytoskeletal organization at interfaces, where cell collective tensions and pressures must be reconciled to achieve tissue integrity.

### Elevated ERK signaling activity in interfacial populations of epithelial cells

We next investigated how signaling may change between the interfaces, since mesenchymal ERK signaling regulates epithelial-mesenchymal boundary integrity and sorting behaviors (Brayford et al., 2019). Specifically, we wondered if increased actin stress fiber density in interior mesenchyme cell collectives and at the interface of interior epithelial tissues correlated with spatial patterns of ERK activation. We therefore fixed and stained for active (phosphorylated) ERK (pERK, **figure 5a**) and quantified the intensity within each tissue and across the interface (**figure 5b**). Radial quantification revealed that pERK levels were highest within the epithelial population in each patterning scheme, with peak intensities in cells located immediately adjacent to the interface and decreasing farther away within the tissue. ERK activity is often spatially anisotropic across developing epithelial tissues, with highest activity at sites of active remodeling, growth, and/or mechanical stress (Hino et al., 2020; Ihermann-Hella et al., 2018; Ishii et al., 2021). Of the two boundary condition types, ERK activation was higher in epithelia that surrounded interior mesenchyme, suggesting that the mesenchymal cell compression observed in **figure 4g** may be associated with strain-induced ERK activation in adjacent epithelial border cells. Our results indicate that the tissue-level context of epithelial-mesenchymal cell contacts modulate ERK activity within the epithelial population. These data reveal engineering control over interfacial signaling relevant to tissue proliferation, motility, and morphogenesis based on cell-biomaterial interface geometry.

**Figure 5.**
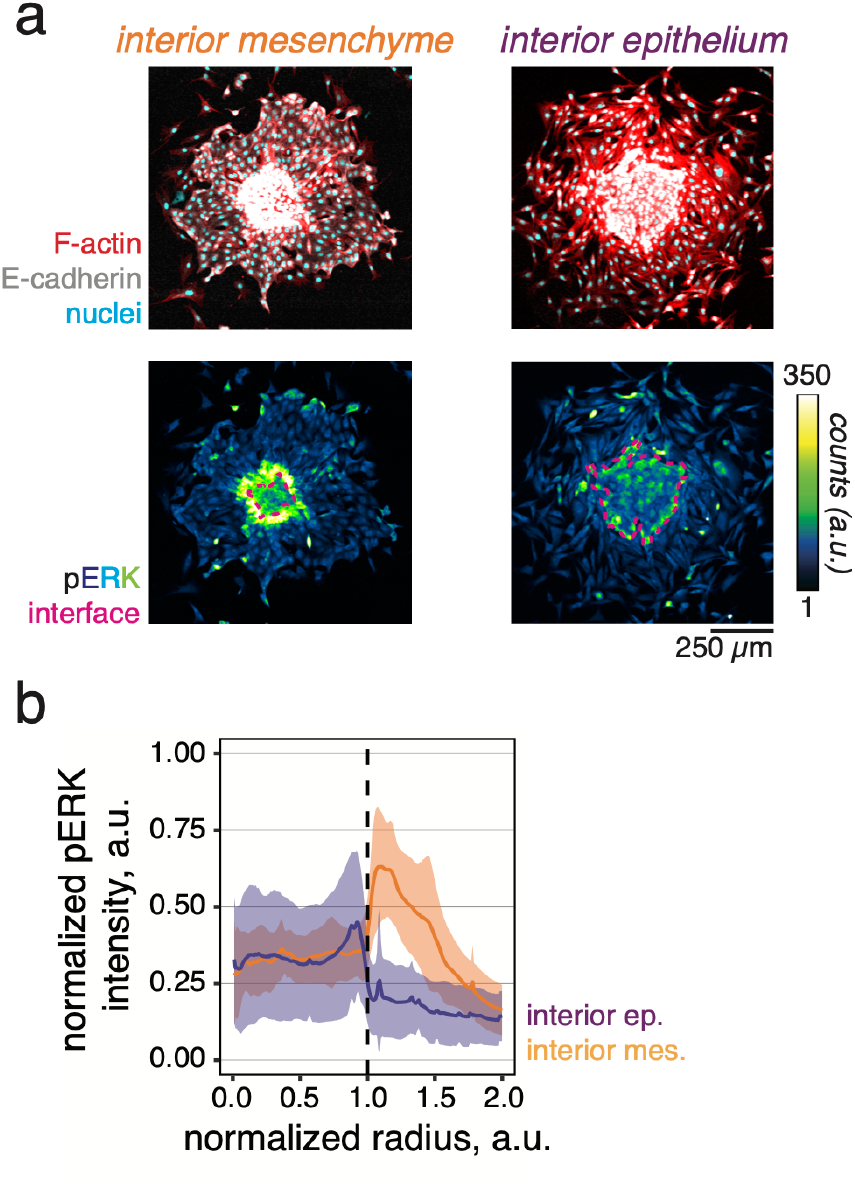
ERK signaling activity in composite tissues depends on interface boundary conditions. **a**. Confocal immunofluorescence for phosphorylated ERK (pERK) and E-cadherin in “interior mesenchyme” and “interior epithelium” tissues and intensity-colored images of pERK staining. **b**. Radial quantification of normalized pERK intensity relative to inner tissue radius for each pattern type. Vertical dashed line marks the tissue interface position (r = 1.0 a.u.), ribbons are mean ± standard deviation (s.d.) of normalized pERK intensity for *n* = 10 interior epithelium and *n* = 15 interior mesenchyme tissues pooled from two independent experiments. Cells were patterned on 7.5%/0.25% (Am/Bis) BP-PA hydrogels sequentially photopatterned with polyT_20_G and polyT_20_F ssDNA (t = 90-120 s exposure at 254 nm) and subsequently functionalized with 20 μg ml^-1^ fibronectin. All tissues were patterned as described in **figure 4** and kept in culture for 12-15 hours to permit interface formation before fixation and staining.

## Discussion

Despite their crucial role in morphogenesis and disease, the cell signaling properties and spatial evolution of tissue interfaces are challenging to study *in vivo* since signaling is often non-cell-autonomous and interface responses can be transient and difficult to analyze in the full 3D context. New engineering approaches to interface construction are required to increase the spatial uniformity, imaging accessibility, and throughput of such studies. Our results extend our previous work (Viola et al., 2020), establishing BP-PA hydrogels as a chassis for orthogonal control of ssDNA-directed cell patterning and cell-ECM adhesion through stepwise photopatterning. This enables both precise design of initial conditions and tracking of adhered interface dynamics for the first time. Importantly, this fabrication process does not significantly alter hydrogel mechanics, cell spreading, or focal adhesion formation, demonstrating that ssDNA patterned hydrogels are suitable for mechanobiology studies. Finally, we identified distinct changes in cellular and cytoskeletal organization and cell signaling at heterotypic cell interfaces as a function of interface geometry alone.

We demonstrate that cell capture can be reliably performed on ∼100 μm-scale features within commercially available clip-on slide chambers, allowing closer adherence to traditional culture protocols. Previous DPAC approaches required specialized, time-intensive and error-prone microfluidic deposition of ssDNA and/or cells (Todhunter et al., 2015; Viola et al., 2020) or iteratively stripping and re-applying photoresist layers onto the substrate between each ssDNA photopatterning step (Scheideler et al., 2020). Here, hundreds to thousands of ssDNA features can be simultaneously deposited across each BP-PA hydrogel using 1-2 minute UV exposure times and ∼20 minutes of subsequent washing steps, allowing a trained user to complete the entire process in ∼30 minutes. Manually aligning and creating a second ssDNA pattern to the first can then be completed within ∼1 hour. Despite vastly simplifying the cell patterning workflow, one tradeoff to our use of commercially available slide chambers is that fluidic control during washing is less precise than previous approaches and potentially exposes cells to higher shear forces during patterning. This can be mitigated by using a microfluidic flow chamber (Scheideler et al., 2020; Todhunter et al., 2015; Viola et al., 2020) to reduce the shear forces from washing and direct patterning down to the single cell scale (∼10 μm feature size). Finally, some cell lines may require additional optimization to improve patterning efficiency or ECM adhesion (**figure S2**).

The DNA patterning approach described here affords a number of experimental handles to tune properties of the engineered tissue microenvironment. Modifying polyacrylamide hydrogel chemistry allows us to tune the elastic hydrogel stiffness and could be extended to incorporate fluorescent fiduciary markers or beads for the measurement of cell-generated traction forces (Pelham and Wang, 1997). Moreover, our fabrication process uses standard, commercially available reagents and UV light sources, with both BPMAC and lipid-ssDNA available by commercial synthesis. We note that BPMAC can also be obtained through straightforward chemical synthesis (Hughes and Herr, 2012). Recent work by Cabral and colleagues also identified commercially available cholesterol-modified ssDNA (Cabral et al., 2021) that is interchangeable with custom lipid-ssDNA used here.

In summary, we have established a new protocol for multiplexed cell patterning on mechanically controlled hydrogel substrates. Access to patterned co-cultures would benefit *in vitro* studies to engineer cell contact-mediated signaling pathways, such as Eph/ephrin (Holland et al., 1996) or planar cell polarity (Strutt and Strutt, 2008). DNA-patterned cell colonies on BP-PA hydrogels could be used in conjunction with synthetic “sender/receiver” stem cell models (Liu et al., 2022) to interrogate the combined roles of mechanics, geometry, and diffusible signaling relays in germ layer patterning. Advanced co-culture models of tumor-stroma interactions (Shen et al., 2014) or cell competition mechanisms (Wagstaff et al., 2016) are other promising applications for this approach.

## Acknowledgements

We thank members of the Hughes lab for helpful discussions. We also thank David Li, Jeff Byfield, and Paul Janmey for microindenter instrument access and training and the staff of the Singh Center for Nanotechnology at the University of Pennsylvania, particularly David Jones, for cleanroom training and assistance with photomask preparation, Arjun Raj for the gift of MDCK II cells, David Odde for LLC-Pk1 cells, Brian Chow for HEK 293T cells, Lukasz Bugaj for 3T3 cells and assistance with lentiviral expression. This work was carried out in part at the Singh Center for Nanotechnology, which is supported by the NSF National Nanotechnology Coordinated Infrastructure Program under grant NNCI-2025608, and at the Penn Cytomics and Cell Sorting Resource Laboratory. This work was supported by NIH grants R35 GM133380 and R01 DK132296 to AJH, T32 HD083185 to JMV, and F32 DK126385 to LSP; an NSF Graduate Research Fellowship to CMP; and the NSF Faculty Early Career Development Program (CAREER grant # 2047271) to AJH.

## Materials & methods

### Oligonucleotides

Photopatterning ssDNA consists of 3’-T_20_X_20_-5’ sequences, where T_20_ is a 20 thymine base sequence and X_20_ is a unique 20 base sequence. We used two previously described sequences (Scheideler et al., 2020; Viola et al., 2020), referred to as polyT_20_F and polyT_20_G. Lignoceric acid (C_23_H_47_COOH, lipid numbers: C24:0) was covalently attached to the universal anchor 3’ end and palmitic acid (CH_3_(CH_2_)_14_COOH, lipid numbers: C16:0) was covalently attached to the universal co-anchor 5’ end through amide linkages (Weber et al., 2014). All oligos and lipid oligos were ordered as custom syntheses (Integrative DNA Technologies), resuspended to either 5 mM (photopatterning ssDNA) or 100 μM (lipid-ssDNA, handles, and fluorescent probes) in DI water and stored at −20ºC. All ssDNA sequences are reported in **table S1**.

### Benzophenone-polyacrylamide gels

1” x 3” glass microscope slides (12-550-A3, ThermoFisher) were cleaned with 0.1% Triton X-100 (T9284, Sigma-Aldrich), rinsed in water, etched in 0.1 N sodium hydroxide (NaOH, S8045, Sigma) for 10 minutes, and dried under compressed air before coating one slide surface with a layer of 3-(trimethoxysilyl)propyl methacrylate (440159, Sigma) and acetic acid in deionized (DI) water (Direct-Q 3 UV water purification system, MilliporeSigma) for 30 minutes to create a surface layer of methacrylate groups that facilitates hydrogel attachment to the slide (Hughes and Herr, 2012). Methacrylate-coated slides were immersed in methanol, washed in DI water, dried under compressed air, and stored for up to 2 weeks before use.

Uniform 30 μm thick hydrogel sheets were cast between the methacrylate-coated glass slide surface and a mechanical grade 4” diameter silicon wafer (University Wafer) containing guide rail shims made from cured SU-8 2025 photoresist (NC9981681, Kayaku Advanced Materials) and rendered hydrophobic by vapor deposition of ∼1 ml dichlorodimethylsilane (DCDMS, 44072, Sigma-Aldrich) *in vacuo* for ∼10 mins. To make wafers, SU-8 was spin coated to 30 μm thickness (SCK-300P, Instras Scientific) onto the wafer surface and exposed to 365 nm UV light (M365LP1, ThorLabs) for 30 s through a custom Mylar mask printed at 20,000 d.p.i. (CAD/Art Services). Excess photoresist was removed using SU-8 developer (NC9901158, Kayaku) followed by alternating washes in isopropanol and acetone. All spin coating, exposure, and heating steps were performed according to manufacturer guidelines. Wafers were washed with alternating 0.1% Triton X-100 and water and dried under compressed air between uses.

BP-PA hydrogel precursor solution contained 3-7.5% acrylamide (Am, 40% w/v stock, 161042, BioRad), 0.035-0.25% N,N-methylenebisacrylamide crosslinker (Bis, 2% w/v stock, 161040, BioRad), 0.06% SDS (5% w/v stock in DI water, 161-0301, Bio-Rad), 0.06% Triton X-100 (5% w/v stock in DI water, T9284, Sigma), and 3 mM N-[3-[(4-benzoylphenyl) formamido]propyl] methacrylamide (BPMAC, custom synthesis, PharmAgra) in 1x Dulbecco’s phosphate buffered saline (DPBS, 10x stock, 14200075, ThermoFisher). First, a partial precursor solution containing Am, Bis, DPBS, and DI water was degassed under vacuum in an ultrasonic cleaning bath (15-337-411, ThermoFisher). SDS, Triton X-100, and BPMAC were then successively added to the precursor with brief vortexing steps between each addition. To initiate polymerization, we successively added ammonium persulfate (0.05% w/v, APS, A3678, Sigma) and N,N,N’,N’-tetramethylenediamine (0.05% v/v, TEMED, T9281, Sigma) and briefly mixed by vortexing. Precursor was injected into the gap between the methacrylate-coated glass slide surface and hydrophobic silicon wafer using a P200 pipette tip and allowed to polymerize for ∼30 minutes. Polymerized hydrogels were rehydrated in 1x DPBS (MT21-031-CV, Corning), carefully lifted off the wafer using a razor blade, and either stored in 1x DPBS at 4ºC overnight or used immediately for photolithography. BPMAC was diluted to 100 mM in DMSO and stored in working aliquots at −20°C. APS was diluted to a 10% w/v stock solution in DI water and stored in working aliquots at −80°C and TEMED was freshly diluted to a 10% v/v stock solution in DI water. Component volumes and base:crosslinker ratios are reported in **table S2**.

### Photomasks

Photomask designs were drawn to scale in LayoutEditor (juspertor) and fabricated in cleanroom facilities by direct writing on 5” x 5” quartz-on-chrome photomasks coated with a layer of IP3500 photoresist (Shipley) using a DWL 66+ laser lithography system (Heidelberg Instruments) equipped with a 10 mm write head. After writing and removing exposed photoresist from features with CD-26 developer (Shipley), washing in DI water, and drying under compressed nitrogen gas, chromium was removed from exposed patches using chromium etchant (Sigma), the surface was washed again with DI water and dried with compressed nitrogen, and excess photoresist was removed by submersion in resist stripper (1165, Shipley) for 3 minutes at 60°C. Photomasks were cleaned with alternating acetone and isopropyl alcohol washes before their first use, then rinsed with 0.1% Triton X-100 and water and dried under compressed air between uses.

### ssDNA photolithography

200-250 μM solutions of polyT_20_F and polyT_20_G ssDNA (5 mM stock at −20°C, **table S1**) in DI water were degassed under vacuum. For simultaneous ssDNA-protein patterning, 200 μM polyT_20_G and either 10 mg ml^-1^ bovine serum albumin (BSA, A2153, Sigma-Aldrich) or 1 μM AlexaFluor 555-labeled BSA (A34786, Fisher) were mixed in 1x DPBS and not degassed. To photopattern ssDNA, BP-PA hydrogels were first completely dried under compressed air, moved to a nitrogen-filled glove box (H50028-2001, Bel-Art), rehydrated with 200-300 μl ssDNA solution, and sandwiched against the photomask region containing the desired pattern. Excess ssDNA solution was removed from the sides by gently wicking with a disposable wipe and the BP-PA gel was exposed through the photomask (λ = 254 nm, I = ∼7 mW cm^-2^, t = 60-120 s) in a UV crosslinker oven (SpectroLinker XL-1000, Spectronics Corporation). Photopatterned BP-PA hydrogels were rehydrated in Tris-acetate-EDTA buffer (1x TAE: 40 mM Tris, 40 mM acetate, 1 mM EDTA, pH 8.0), gently lifted off the photomask using a razor blade, and then washed in 1x TAE + 0.1% SDS followed by 1x TAE and stored in 1x DPBS.

To register and align multiple ssDNA patterns on the same gel, we incorporated a previously described “cross and window” alignment scheme into the photomask design (Viola et al., 2020). Briefly, the hydrogels were dried under compressed air and “cross” features were labeled using a drop of 1 μM of the appropriate 5’-6-fluorescein amidite (FAM) probe (e.g., FAM_G’, **table S1**) for ∼5 minutes, followed by washing in 1x TAE + 0.1% SDS and 1x TAE. Hydrogels were then incubated in the second ssDNA sequence, assembled against the photomask in a nitrogen glove box, and then manually aligned to the photomask “window” regions using the GFP filter set on a benchtop epifluorescence microscope. The second ssDNA patterns were UV-exposed, washed, and stored as described above.

### Fibronectin functionalization through NHS esters

To derivatize acrylamide chains with bioreactive NHS esters, a solution of 0.01% w/v N,N-methylenebisacrylamide crosslinker (stored as 0.2% w/v stock in DI water) buffered with 50 mM HEPES (4-(2-hydroxyethyl)-1-piperazineethanesulfonic acid, 0.5 M stock at pH 6.0, 40820004, bioWORLD) and was degassed under vacuum in an ultrasonic bath. Next, the photoinitiator LAP (0.09% w/v, lithium phenyl-2,4,6-trimethylbenzoylphosphinate, 900889, Sigma) and N2 acrylic acid (0.1% w/v, acrylic acid-N-hydroxysuccinimide ester, A8060, Sigma) were successively added and mixed by vortexing. LAP stock solutions (0.9% w/v in DI water) were stored at 4°C for up to 2 weeks. N2 was stored at −20°C and freshly prepared (0.3% w/v in 50% ethanol/50% DI water).

Hydrogels were dried and rehydrated in the NHS mixture, drawing the liquid evenly across the hydrogel surface using a DCDMS-treated glass slide. Hydrogels were sandwiched between the supporting slide and the DCDMS-treated slide and UV-exposed (λ = 365 nm, I = 15 mW cm^-2^) for 20 minutes, then rehydrated in ice cold DI water. We gently lifted off the DCDMS-treated slide and washed twice in ice cold DI water containing 100 μM sodium chloride (NaCl, S6191, Sigma). Hydrogels were then partially dried and a plastic clip-on 8-well slide with a silicone gasket (CCS-8, MatTek) was assembled around the region containing photopatterned ssDNA features and wells were incubated in 10-20 μg ml^-1^ bovine serum fibronectin (1 mg ml^-1^, F1141, Sigma) diluted in sterile 50 mM HEPES (pH 8.5) overnight at 4°C. Next day, fibronectin solution was removed and wells were washed in 50 mM HEPES (pH 8.5) containing 100 mM glycine (G8898, Sigma) for 1 hour at room temperature, followed by 3-4 washes in sterile 1x DPBS.

Sulfo-SANPAH (sulfosuccinimidyl 6-(4’-azido-2’-nitrophenylamino)hexanoate) was used as an alternative approach to derivatize hydrogels with NHS esters for ECM conjugation (**figure S2**). After drying the photopatterned hydrogel and assembling the clip-on slide, hydrogels were rehydrated with 1 mg ml^-1^ Sulfo-SANPAH diluted in DI water (22589, ThermoFisher) and irradiated with collimated UV light (λ = 365 nm, I = 15 mW cm^-2^) for 10 minutes. Exposed gel surfaces were washed in 1x DPBS followed by two washes in 50 mM HEPES (pH 8.5) and incubated overnight in 10-20 μg ml^-1^ fibronectin in 50 mM HEPES (pH 8.5) at 4°C before washing and adding cells. Sulfo-SANPAH aliquots (1 mg in 20 μl DMSO) were flash frozen in liquid nitrogen and stored at −80°C.

### Cell culture

MDCK II epithelial cells were cultured in minimum essential medium (MEM, Earle’s salts and L-glutamine, MT10-010-CV, Corning) supplemented with 10% fetal bovine serum (FBS, MT35-010-CV, Corning) and 1x pen/strep (100 IU ml^-1^ penicillin, and 100 μg ml^-1^ streptomycin, 100x stock, 15140122, Invitrogen) and passaged with 0.25% trypsin-EDTA (2530056, Corning). 3T3 mouse embryonic fibroblasts were maintained in Dulbecco’s minimum essential medium (DMEM, 4.5 g l^-1^ glucose, L-glutamine, and sodium pyruvate, MT10-013-CV, Corning) supplemented with 10% calf serum (SH3008703, HyClone) and 1x pen/strep. LLC-Pk1 epithelial cells were cultured in MEM supplemented with 10% FBS, 1 mM sodium pyruvate (100 mM stock, 11360070, Invitrogen), and 1x pen/strep and passaged with 0.25% trypsin-EDTA. Human embryonic kidney 293T (HEK 293T) cells were cultured in DMEM supplemented with 10% FBS and 1x pen/strep and passaged with 0.05% trypsin-EDTA. MDCK H2B-Venus cell lines are previously described (Viola et al., 2020). All cells were cultured at 37ºC and 5% CO_2_ in polystyrene 75 cm^2^ or 182 cm^2^ flasks kept in a humidified incubator..

### Lentiviral transduction

Cell lines expressing H2B-infrared fluorescent protein 670 (H2B-iRFP) were established by lentiviral transduction. To generate H2B-iRFP lentiviral particles HEK 293T cells were plated in a 6-well plate (7×10^5^ cells per well) and transiently transfected with 1.5 μg of pLenti-pGK-DEST-H2B-iRFP vector (Addgene # 90237, RRID: Addgene_90237, gift from Markus Covert), 1.33 μg pCMVd8.1 (Addgene #12263, RRID: Addgene_12263, gift from Didier Trono), and 0.17 μg pMD2.G (Addgene #12259, RRID: Addgene_112854, gift from Didier Trono) using the calcium phosphate method (Bugaj et al., 2015). After 48 hours, particles were collected from the supernatant, centrifuged for 3 minutes at 800xg, and sterile filtered through a 0.45 μm syringe filter to remove cell debris. Filtered particles were either immediately introduced to cells (10^5^ target cells plated in a 6-well plate) in antibiotic-free culture medium supplemented with 8 μg ml^-1^ polybrene (TR-1003-G, EMD Millipore) or stored in single use aliquots at −80ºC. After ∼48 hours, transduced cell lines were expanded and enriched for fluorescent expression using a BD Influx cell sorter (BD Biosciences).

### Lipid-ssDNA cell labeling and patterning

For a typical patterning experiment (1-2 BP-PA gels) we prepared a 70-80% confluent T182 flask (∼1.0-2.0×10^7^ cells) for each cell line. Cells were dissociated from flasks with trypsin and (if appropriate for the experiment) labeled with 1 μm CellTracker Deep Red (C34565, Fisher) cytoplasmic dye in serum-free DMEM for 15-30 minutes at 37°C prior to lipid-ssDNA labeling. Suspended cells were pelleted and washed twice in 1x DPBS (Ca^2+^/Mg^2+^-free, MT21-031-CV, Corning), transferred to a 2 ml Eppendorf tube, and resuspended in 100 μl DPBS. Lipid anchor, lipid co-anchor, and handle ssDNA (5 μl each, **table S1**) were successively added with 10 minute incubations between each. Labeled cells were washed twice and resuspended in 0.5-1 ml DPBS.

To introduce cells, we first aspirated the liquid from each well and added ∼100 μl of labeled cell suspension. Cells were allowed to settle for ∼5 minutes before excess cells were removed by gently washing with 1x DPBS using a P200 pipette tip. Excess cells could be returned to the Eppendorf tube and stored on ice to be reintroduced to a new substrate. Next, the hydrogel surface was gently washed in DPBS using a P200 tip or vacuum aspirator to remove liquid.

Alternatively, the 8-well slide could be disassembled and labeled cells introduced to the ssDNA deposits across the entire hydrogel surface. After allowing cells to settle on the ssDNA patterns, we iteratively washed the hydrogel surface by immersion in 1x DPBS to remove non-adherent cells. After the last wash, the remaining DPBS was gently removed from wells and replaced with a complete imaging medium. Imaging medium consisted of phenol-free DMEM (4.5 g l^-1^ glucose, L-glutamine, and 25 mM HEPES, 21063-029, Invitrogen) supplemented with 10% FBS, 1 mM sodium pyruvate, and 1x pen/strep). To visualize ssDNA patches during time lapse imaging (**figure 1**) we stained ssDNA with 1-5x SYBR Gold nucleic acid gel stain (S11494, ThermoFisher) in 1x DPBS for 5-10 minutes and washed twice in 1x DPBS before introducing culture medium. SYBR Gold was resuspended to a 10,000x stock in DMSO and stored in working aliquots at −20ºC.

### Microindentation

We adapted a previously described micro-indenter device and approach to measure *E* of BP-PA hydrogels (Levental et al., 2010). Hydrogels were cast on methacrylate-coated glass slides at various Am/Bis ratios (**table S2**) with 5 μl far red fluorescent microspheres (F8807, ThermoFisher) added to the prepolymer mixture. To cast hydrogels, ∼250 μL of pre-polymer mixture was introduced into a gap between the methacrylate-treated glass slide and a DCDMS-treated hydrophobic 2” x 3” slide (2947, Corning) created using double layered laboratory tape shims as guides. Following polymerization and rehydration in 1x DPBS, we uniformly photopatterned half of each hydrogel with 200 μM polyT_20_G ssDNA (λ = 254 nm, I = 7 mW cm^-2^, t = 90 s) through a 1 mm thick quartz microscope slide (26011, Ted Pella) using aluminum foil to mask the other side from UV exposure.

The microindentation apparatus consisted of a 255 μm diameter blunt end indenter fabricated from cylindrical 30 gauge (AWG) SAE 316L stainless steel wire attached to a tensiometric sensor and stepper motor (L4018S1204-M6, Nanotec) to control indenter z position. After calibrating the spring constant of the sensor on a rigid (glass) surface, we indented BP-PA hydrogels using an approach rate of 12.5 μm s^-1^ while simultaneously recording time, force, and indenter z-position. *E* was calculated from the linear part of force versus indentation depth curves by the following relationship (**eqn. 1**), assuming a soft homogeneous material of finite thickness and a rigid, cylindrical indenter:

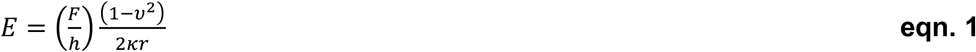

In **eqn. 1** 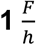 is the slope of the F-d curve, *ν* is the Poisson ratio (0.457 for polyacrylamide hydrogels (Takigawa et al., 1996)), *κ* is the Hayes correction factor for finite sample thickness (Hayes et al., 1972), and 2*r* is the probe diameter (255 μm). Values for *κ* were estimated using hydrogel thickness (t) measurements obtained from confocal z-stacks (10 μm interval) of fluorescent microspheres embedded within the gel. F-d curves were fit to a linear model using the Matlab Optimization and Curve Fitting toolbox.

### Imaging

In experiments quantifying cell capture (**figure 1c**) and fibroblast spreading experiments (**figure 2c-d**), patterned hydrogels were imaged using a Nikon Ti2 epifluorescence microscope (Nikon Instruments) equipped with a motorized stage, CMOS camera (DS-Qi2, Nikon), LED transmitted and epifluorescence illumination (Sola II light engine, Lumencor), single-pass DAPI, FITC, TRITC, and Cy5 filter sets (Chroma), 4x/0.25 numerical aperture (NA), 10x/0.25 NA, and 20x/0.75 NA lenses. For other live and immunofluorescence imaging, we used a Ti2 microscope equipped with a CSU-W1 spinning disk (Yokugawa), a solid state laser launch (100 mW 405, 488, and 561 nm lasers and a 75 mW 640 nm laser), a white light LED for transmitted illumination, a motorized stage, a Prime 95B back-thinned CMOS camera (Teledyne Photometrics), and 4x/0.25 NA, 10x/0.25 NA, and 20x/0.45 NA lenses. A sealed enclosure built around the microscope stage (OkoLab) provided stable environmental conditions at 37°C and 5% CO_2_ for the duration of the experiment. Microscopes were under control of NIS Elements AR software (version AR 5.11.00)

### Immunofluorescence

To visualize fibronectin coupled to BP-PA gels alongside photopatterned polyT_20_F ssDNA (**figure 1e**) we first incubated the gels in a rabbit anti-fibronectin antibody (1:100, ab23750, Abcam, RRID: AB_447655) diluted in 1x DPBS for 45 minutes followed by donkey anti-rabbit IgG-AlexaFluor 647 (1:200, A31573, ThermoFisher, RRID: AB_2536183) and 1 μM FAM_F’ probe (**table S1**) diluted in 1x DPBS for 30 minutes. Stained gels were washed 1x DPBS before imaging.

Cells were fixed in 4% paraformaldehyde (PFA, 16% stock, 15710, Electron Microscopy Sciences) diluted in 1x DPBS for 10 minutes, permeabilized in 0.5% v/v Triton X-100 (10 minutes) in 1x DPBS, and blocked with 1% BSA in PBS-T (1x DPBS + 0.1% v/v Tween-20, #9480, EMD Millipore) for 30-60 minutes. All fixation, permeabilization, and blocking steps were performed at room temperature. Fixed and blocked cells were incubated in primary antibodies for 2 hours at room temperature or 4°C overnight and secondary antibodies for 1 hour at room temperature with three PBS-T washes following each incubation. Primary antibodies were diluted in blocking buffer: mouse anti-vinculin (7F9) (1:100, #14-9777-82, eBiosceince, RRID: AB_2573028), mouse anti-E-cadherin (clone 34, 1:200, 610404, BD Biosciences, RRID: AB_397787), rabbit anti-phospho-p44/42 MAPK (ERK1/2) (1:200, #4370, Cell Signaling Technologies, RRID: AB_2315112), and rabbit anti-E-cadherin (24E10) (1:200, #3195, Cell Signaling Technologies, RRID: AB_2291471). Secondary antibodies (all raised in donkey) were used at 1:1000 dilution: anti-rabbit AlexaFluor 488 (A21206, Invitrogen, RRID: AB_2535792), anti-mouse AlexaFluor 555 (A-31572, ThermoFisher, RRID: AB_162543), anti-rabbit AlexaFluor 555 (A31572, Invitrogen, RRID: AB_162543), and anti-rabbit AlexaFluor 647 (A31573, ThermoFisher, RRID: AB_2536183). Nuclei and F-actin were counterstained with 4′,6-diamidino-2-phenylindole dihydrochloride (DAPI, 300 nM, D1306, ThermoFisher) and phalloidin-AlexaFluor 647 (1:100, A22287, Invitrogen) added along with the secondary antibodies. For fibroblast spreading assays (**figure 2c-d**), we fixed and permeabilized as described and then directly stained cells for 1 hour with phalloidin-AlexaFluor 647, DAPI, and 1 μM FAM_G’ probe diluted in PBS-T without blocking.

### Image analysis

All image analysis was performed using ImageJ/Fiji (Schindelin et al., 2012). Cells captured on ssDNA patches in **figure 1** were manually counted from fluorescence signal (CellTracker dye) or brightfield images using the *multi point* selection tool in ImageJ/Fiji. Patch boundaries were determined using the SYBR Gold fluorescence signal. To count multiple cell populations on ssDNA patterns that were not labeled (chess board in **figure 3** and **figure S3**) we drew a 250 μm square region of interest (ROI) around the middle of each captured cell cluster and counted cells within the bounding region.

To measure fibroblast spread areas (**figure 2c-d**) we acquired image stacks at 20x magnification on the Ti2 widefield system and manually thresholded the phalloidin-AlexaFluor 647 channel in ImageJ/Fiji to obtain a binarized image containing cell outlines. Next, we used the *erode, fill holes*, and *analyze particles* functions with a minimum area (excluding objects < 500 μm^2^) to obtain ROIs for individual cells. We excluded cells on the edge of the image and limited our analysis to single cells by using the nucleus (DAPI) channel to exclude ROIs that contained more than one nucleus. To obtain images for focal adhesion measurements (**figure 2e-f**) we acquired z-stacks (13 frames, 2.5 μm z-step size, 30 μm total) with the 20x lens using a 1.5x intermediate zoom lens (30x objective magnification). We measured focal adhesion lengths from the vinculin immunofluorescence channel manually using the *line segment* tool in Fiji/ImageJ. In both sets of experiments, we used the FAM_F’ or FAM_G’ signal to discriminate between UV-exposed and unexposed control hydrogel regions.

We employed a custom Fiji/imageJ radial analysis plugin (Dobretsov and Romanovsky, 2006) to quantify fluorescence intensity in composite epithelial-mesenchymal tissues (**figure 4-5**). We used the *draw polygon* function to trace the boundary between the inner and outer tissues and radially transformed the ROI to obtain a 200 pixel radius circular ROI with the interface located at r = 100 pixels (normalized to r = 1.0 a.u.). The use of a radial transformation enables an “apples-to-apples” comparison of irregularly shaped tissues or interfaces. After removing background by subtracting a fixed value, fluorescence intensities of F-actin, E-cadherin, or pERK were calculated at each evenly spaced radial bin (pixels in **figure S4**) and normalized to the maximum pixel intensity across all measurements. We excluded tissues with an incomplete border (e.g., where <60% of the circumference was surrounded by the second cell population) from our analysis.

### Statistical analysis and reproducibility

Statistics and plotting were performed using R (version 4.2.0, R Core Team) running on RStudio version 2022.02.0. Exact numbers of measurements (*n*), numbers and types of independent replicates, and statistical tests used to compare results and compute *p*-values are indicated in the figure legends. Two-sided *p*-values are reported unless otherwise noted. We used Welch’s two-sided t-test to facilitate comparisons between groups where data were normally distributed (as determined by the Shapiro-Wilk non-normality test with a *p* = 0.05 cutoff) and non-parametric tests otherwise. For microindentation measurements of *E* (**figure 2b**) we used paired t-tests to compare mean elastic modulus of exposed (*E*_*uv*_) and unexposed (*E*_*ctrl*_) regions of the same hydrogel. *E* for each hydrogel is the mean of 3 measurements per side for a total of 6 measurements per hydrogel; overall mean ± s.d. of *E* for *n* hydrogels of each composition is described in **table S2**. We used a two-tailed Kolmogorov-Smirnov test to compare cumulative distributions of fibroblast spreading (**figure 2d**) or focal adhesion lengths (*L*_*adhesion*_) on unexposed and photopatterned hydrogel regions (**figure 2f**). All data are representative of at least two independent biological replicate experiments, where appropriate.

