## Supplemental information for "Independent control over cell patterning and adhesion on hydrogel substrates for tissue interface mechanobiology"

**Authors & affiliations**

Louis S. Prahl[1], Catherine M. Porter[1], Jiageng Liu[1], John M. Viola[1], & Alex J. Hughes[1,2,3,*]
[1]Department of Bioengineering, [2]Department of Cell & Developmental Biology, and [3]Institute for Regenerative Medicine, University of Pennsylvania, Philadelphia, PA, USA 19104

### Supplemental notes

*Supplemental Note 1: Comparison of NHS derivatization methods for BP-PA hydrogels*
In the present study, we attempted several different methods of ECM coupling chemistry. UV irradiation (λ = 254 nm) of BPMAC across the entire hydrogel surface in the presence of soluble fibronectin failed to produce subsequent cell attachment (**figure S2a**). When we used Sulfo-SANPAH as our NHS source, we found that MDCK cells could adhere to the unexposed hydrogel surface but cleared from regions where BPMAC had been exposed (**figure S2b**). This occurred even in cases where ssDNA had not been present in the photopatterning solution (data not shown) but could be overcome by adding bovine serum albumin (BSA) or fibronectin to the ssDNA patterning mixture. This dramatically improved the adhesion of MDCK cells, yet other cell types typically remained poorly adherent to UV-exposed hydrogel regions (**figure S2c**). Changes in polyacrylamide hydrogel base:crosslinker ratio reportedly cause variability in Sulfo-SANPAH activation and protein coupling (Kandow et al., 2007). We suspect that BPMAC exposure to deep UV (λ = 254 nm) may destroy Sulfo-SANPAH binding sites or create hydrophobic regions where cells have difficulties accessing ECM ligands. We think it is unlikely that changes in crosslinking are responsible, given that microindentation revealed no differences in *E*, except in the softest hydrogel mixtures (3%/0.05% Am/Bis hydrogels, see: **figure 3b**). Given that Sulfo-SANPAH performed poorly compared to the NHS-acrylic acid method (see: **materials & methods**), we adopted the latter for experiments in the present study.

### Supplemental figures

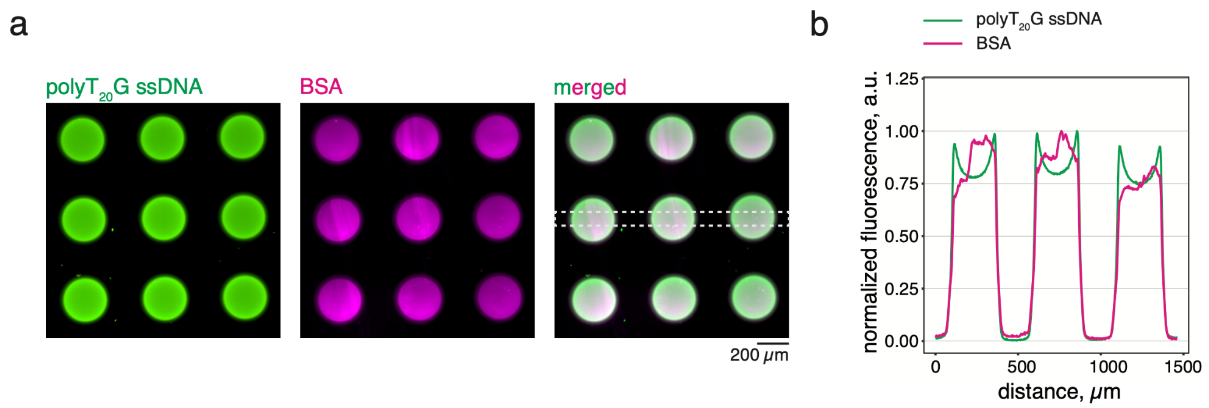


**Figure S1. Simultaneous patterning of ssDNA and proteins by UV-exposure of BPMAC co-monomers. a.** Image of polyT$_{20}$G ssDNA and BSA-AlexaFluor 555 photo patterned (**I**$_{254\ nm}$ = 7 mW cm$^{-2}$, t = 90 s) into arrays of 250 μm diameter circular features. **b.** Line-scan across the dashed 25-pixel wide region in showing fluorescence traces from hydrogel-bound ssDNA and BSA. Each channel is normalized to maximum pixel intensity. polyT$_{20}$G is visualized using 1 μM FAM_G' probe (**table S1**).

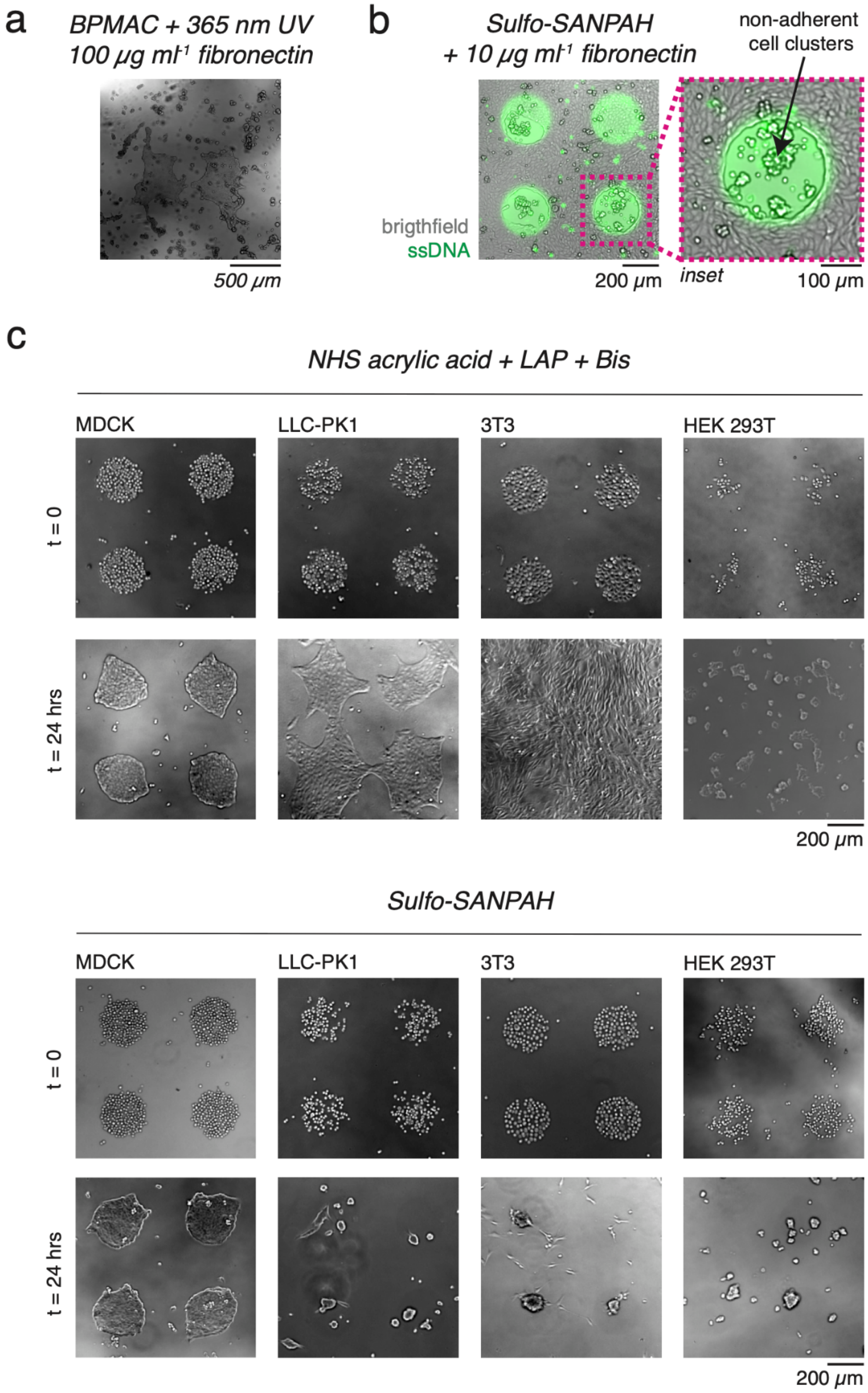


**Figure S2. Comparison of DNA-mediated cell capture and ECM-mediated cell adhesion for various functionalization techniques. a.** MDCK cells plated on a BP-PA hydrogel surface irradiated with UV light ($I_{365\ nm}$ = 15 mW cm$^{-2}$, t = 10 minutes) in the presence of 100 μg ml$^{-1}$ fibronectin and allowed to adhere for 16 hours. Cells did not evenly adhere across the entire surface, while some areas supported adhesion, others showed cell detachment and aggregation into floating clusters. **b.** MDCK cells introduced to a BP-PA hydrogel surface bearing arrays of 250 μm diameter circular ssDNA features frequently failed to invade onto the feature surface. Cells atop the ssDNA feature show evidence of de-wetting and clustering (*inset*). polyT$_{20}$F ssDNA was photopatterned for 120 seconds ($I_{254\ nm}$ = 7 mW cm$^{-2}$) in a solution of 1x DPBS that also contained 100 μg ml$^{-1}$ fibronectin. In a subsequent step, fibronectin (10 μg ml$^{-1}$ in 50 mM HEPES, pH 8.5) was conjugated to the entire hydrogel surface following UV activation of Sulfo SANPAH ($I_{365\ nm}$ = 15 mW cm$^{-2}$, t = 10 minutes). MDCK cells were added in two steps: 1) a lipid-ssDNA labeled population was first introduced to ssDNA features, exchanged for complete culture medium, and allowed to adhere for ~1 hour before adding 2) a solution of 10$^5$ MDCK cells in complete culture medium was layered on top of the patterned cells. Images were acquired 16 hours after the addition of the first group. ssDNA features were subsequently labeled with 5x SYBR Gold nucleic acid stain before imaging. **c**. Comparison of cell adhesion effectiveness for four different DNA-patterned cell lines and two surface functionalization techniques on BP-PA hydrogels. *Top*, functionalization via LAP, bis-acrylamide, and N2 acrylic acid (Lakins et al., 2012). *Bottom*, functionalization via Sulfo-SANPAH. In both cases, 20 μg ml$^{-1}$ fibronectin was coupled to the surface via overnight incubation. ssDNA (polyT$_{20}$G) was photopatterned ($I_{254\ nm}$ = 7 mW cm$^{-2}$, t = 90 s) in a solution of 1x DPBS containing 10 mg ml$^{-1}$ BSA and lipid-ssDNA labeled cells were introduced with the handle G' ssDNA. See: **materials and methods**.

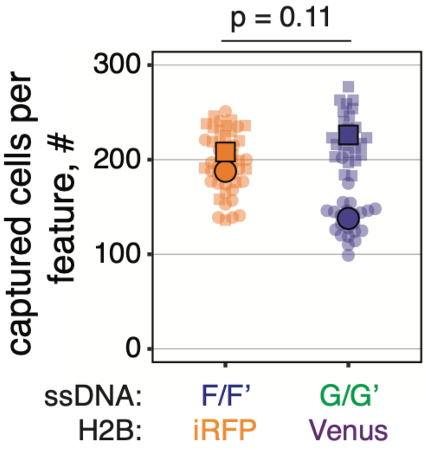


**FIgure S3. Quantification of cell capture for two populations of MDCK cells in a chess board pattern.** MDCK cells expressing either H2B-iRFP or H2B-Venus captured on arrays of 250 µm square ssDNA deposits using the F/F' or G/G' ssDNA pairs, $n$ = 24 randomly selected features per ssDNA sequence pooled from two wells (technical replicates) each from two independent experiments. Experiment means (black borders) are overlaid onto individual measurements, $p$-value was obtained using a two-sided Wilcoxon rank sum test. See also: **figure 3**.

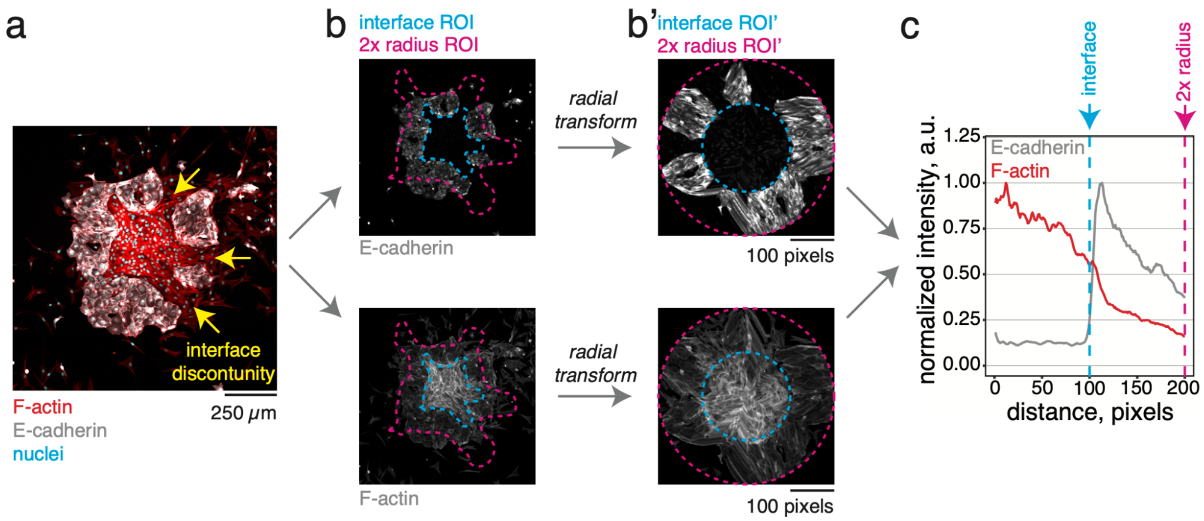


**Figure S4. Radial quantification of fluorescent markers across patterned interfaces. a.** Example image of a composite tissue containing MDCK epithelial cells (marked by E-cadherin and actin) and 3T3 fibroblasts (only actin) in the "mesenchyme inside" format. Nuclei are counter-stained with DAPI. Yellow arrows show locations of discontinuities within the bounding epithelium. **b.** Single channel images of F-actin and E-cadherin before and after radial transformation of the interface ROI (cyan dashed line). Magenta dashed line represents the dilation of the ROI out to 2x radius. **b'.** Images following radial transformation (Dobretsov and Romanovsky, 2006). ROI' are the radially transformed ROIs shown in panel **b**, note the conversion to pixel units. **c.** Fluorescence intensity traces of E-cadherin and F-actin signal normalized to the brightest pixel bin in each channel. Dashed lines show the interface position (cyan) and 2x radius (magenta). See also: **figures 4-5** and **figure S5**.

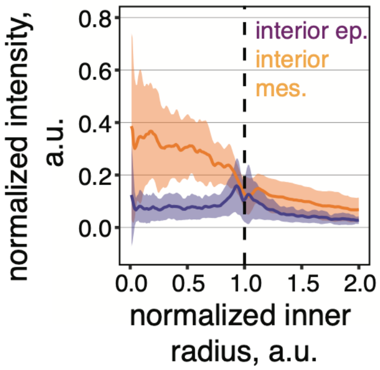


**Figure S5. Radial quantification of nuclear signal as an approximation of cell density.** Radial quantification of nuclear (DAPI) signal intensity for the immunostained tissues in **figure 4**. Ribbons are mean ± s.d. of $n$ = 15 interior epithelium and $n$ = 11 interior mesenchyme tissues collected from two independent experiments. Individual traces were normalized to the maximum pixel intensity in a given experiment.

**Supplemental tables**

| ssDNA sequence | name |
|---|---|
| *patterning ssDNA* | |
| 5'-TTTTTTTTTTTTTTTTTTTTTAGAAGAAGAACGAAGAAGAA-3' | polyT$_{20}$F |
| 5'-TTTTTTTTTTTTTTTTTTTTTAGCCAGAGAGAGAGAGAGAG-3' | polyT$_{20}$G |
| *lipid-ssDNA anchors* | |
| 5'-TGGAATTCTCGGGTGCCAAGGGTAACGATCCAGCTGTCACT-lignoceric-amide-3' | universal anchor |
| 5'-palmitic-amide-AGTGACAGCTGGATCGTTAC-3' | universal co-anchor |
| *handle ssDNA* | |
| 5'-CCTTGGCACCCGAGAATTCCATTTTTTTTTTTTTTTTTTTTTCTTCTTCGTTCTTCTTCT-3' | F' handle |
| 5'-CCTTGGCACCCGAGAATTCCATTTTTTTTTTTTTTTTTTTTCTCTCTCTCTCTCTCTGGCT-3' | G' handle |
| *fluorescent probes* | |
| 5'-5(6)-FAM-TTTCTTCTTCGTTCTTCTTCT-3' | FAM_F' |
| 5'-5(6)-FAM-CTCTCTCTCTCTCTCTGGCT-3' | FAM_G' |
| 5'-Cy5-TTTCTTCTTCGTTCTTCTTCT-3' | Cy5_F' |

**Table S1. ssDNA oligomers used for photopatterning, cell anchoring, and fluorescent labeling.** 5(6)-FAM = 5(6)-Carboxyfluorescein dye.

| Am/Bis ratio | 40% Am | 2% Bis | DI $H_2O$ | 10x DPBS | 5% SDS | 5% Triton -X100 | 100 mM BPMAC | 10% APS | 10% TEMED | $E_{ctrl}$, kPa (*n* of gels) | $E_{uv}$, kPa |
|---|---|---|---|---|---|---|---|---|---|---|---|
| 3%/0.05% | 37 µl | 13 µl | 372 µl | 50 µl | 3 µl | 3 µl | 15 µl | 3 µl | 3 µl | 2.4 ± 0.6 (*n* = 4) | 2.8 ± 0.7 |
| 3%/0.1% | 37 µl | 25 µl | 360 µl | 50 µl | 3 µl | 3 µl | 15 µl | 3 µl | 3 µl | 8.7 ± 6.3 (*n* = 2) | 8.8 ± 4.9 |
| 7.5%/0.035% | 94 µl | 9 µl | 320 µl | 50 µl | 3 µl | 3 µl | 15 µl | 3 µl | 3 µl | 17.3 ± 5.9 (*n* = 4) | 16.4 ± 5.5 |
| 7.5%/0.07% | 94 µl | 18 µl | 311 µl | 50 µl | 3 µl | 3 µl | 15 µl | 3 µl | 3 µl | 24.1 ± 5.6 (*n* = 3) | 23.5 ± 3.9 |
| 7.5%/0.25% | 94 µl | 68 µl | 261 µl | 50 µl | 3 µl | 3 µl | 15 µl | 3 µl | 3 µl | 36.3 ± 7.6 (*n* = 4) | 31.5 ± 19.2 |

**Table S2. BP-PA hydrogel prepolymer formulations and summary of elastic modulus measurements.** Am = acrylamide, Bis = N,N-methylenebisacrylamide crosslinker, DPBS = Dulbecco's phosphate buffered saline, SDS = sodium dodecyl sulfate, BPMAC = N-[3-[(4-benzoylphenyl) formamido]propyl] methacrylamide, APS = ammonium persulfate, TEMED = N,N,N',N'-tetramethylenediamine. Volumes are calculated for a 500 µl total precursor stock. Units in parentheses on the top row represent the stock solution concentration. $E_{ctrl}$ is the bulk elastic modulus for unexposed regions and $E_{uv}$ is the bulk elastic modulus for UV-exposed regions; values of $E_{ctrl}$ and $E_{uv}$ are the mean of three measurements per unexposed and UV-exposed side of each hydrogel, respectively. Measurements represent mean ± s.d. for *n* gels. Mixtures used to cast BP-PA hydrogels for microindentation measurements replaced 5 µl of the DI $H_2O$ volume with 5 µl deep red fluorescent microspheres.

### Supplemental movies

**Movie S1. Epithelial microtissues spreading on ssDNA patterns of fixed area and variable shape.** Time lapse image sequence of MDCKs expressing H2B-iRFP (magenta) adhering to circular, triangular, square, and star shaped ssDNA features labeled with SYBR Gold (green). Patterns were designed with a fixed area equivalent to a 200 μm diameter circle ($A = 3.14 \times 10^4$ $\mu m^2$). Images were acquired every 20 minutes for 10 hours, starting 1 hour after DNA patterning. See also: **figure 1e**.

**Movie S2. Shape change of composite tissues consisting of a circle of mesenchymal cells surrounded by an epithelial ring.** Time lapse image sequence of MDCKs expressing H2B-Venus (magenta) surrounded by a ring of 3T3s expressing H2B-iRFP (yellow). Patterns consist of an inner circle (r = 175 μm) surrounded by an annulus (r = 75 μm), each with a nominal area of A = 97264 $\mu m^2$. Images were acquired every 2 hours for 10 hours total, starting 1 hour after DNA patterning. See also: **figure 4b**.

**Movie S3. Shape change of composite tissues consisting of a circle of epithelial cells surrounded by a mesenchymal ring.** Time lapse image sequence of 3T3s expressing H2B-iRFP (yellow) surrounded by a ring of MDCKs expressing H2B-Venus (magenta). Patterns consist of an inner circle (r = 175 μm) surrounded by an annulus (r = 75 μm), each with a nominal area of A = 97264 $\mu m^2$. Images were acquired every 2 hours for 10 hours total, starting 1 hour after DNA patterning. See also: **figure 4b**.
